# Spinotrode: long-term intraspinal electrophysiological recordings to unravel dorsal horn neuron dynamics in behaving mice

**DOI:** 10.64898/2026.08.19.745481

**Authors:** Juliette Viellard, Louison Brochoire, Michelle Janusz, Christopher Dedek, Franck Aby, Rabia Bouali-Benazzouz, Feng Wang, Benoit Gosselin, Steven A Prescott, Abdelhamid Benazzouz, Yves De Koninck, Pascal Fossat

## Abstract

Recording single-unit neural activity in the spinal cord in freely moving rodents is crucial for understanding spinal network dynamics but remains challenging due to specific biomechanical constraints. So far, the vast majority of studies have been conducted in anesthetized or restrained animals, limiting the correlation of neuronal activity with naturalistic behaviours. Here we introduce the Spinotrode, a vertebral implant able to stably record signals (over several weeks) at the single-cell level in the bilateral dorsal horns of adult mice. It is engineered to minimize postural constraints and interrogate spinal activity during sensory stimulation and motor behaviours. No functional impairment or tissue damage was apparent. Spinotrode recordings allowed identification of distinct functional types of neurons associated with paw withdrawal, revealed dorsal horn activity during locomotor behaviour distinct from proprioception and touch, and uncovered contralateral sensory activation upon nociceptive reflex responses, which forces reassessment of data obtained in anesthetized animals.

## Introduction

Sensorimotor integration emerges from continuous interactions between the peripheral and central nervous systems, with the spinal cord playing a critical role as a central hub. It relays sensory inputs to supraspinal regions^1–3^ while integrating descending signals to regulate sensory transmission^4–6^ and initiate motor outputs^7^. Classical *ex vivo* and anesthetized *in vivo* electrophysiological approaches have provided insights into the functional properties of individual neurons within spinal segments^8–11^. However, these approaches disrupt network integrity and profoundly alter neuronal firing dynamics^12–14^, restricting analyses to evoked responses. Recordings from freely moving animals – a landmark long achieved in brain regions^15–17^ – provide a solution to bridge *in vivo* neuronal activity and complex, unconstrained behaviour. High-density electrode devices commonly used in the brain ^18–20^ have been difficult to use in the spinal cord due to its small size, limited accessibility, and movements associated with breathing and locomotion^21^. Despite numerous attempts ^22–31^, stable, long-term and accurate spinal recordings have not been achieved without interfering with natural posture and locomotion. Most of these studies are limited to low-resolution multiunit or field potential recordings^22–25^, short-term measurements^26,27,30,32,33^, or rely on heavy and unwieldy implants^31^.

To overcome these limitations, we developed a novel electrode system designed for bilateral spinal recordings in freely moving mice, enabling chronic monitoring of neural activity while preserving natural behaviour, without measurable interference with motor, immune, nor sensory functions. Using this device, we revealed the patterns of neuronal firing during spontaneous movements and in response to peripheral stimulation, identifying mechanosensitive, thermosensitive, and polymodal units with distinct coding dynamics associated with nociceptive and/or non-nociceptive input. Spike waveform analysis further revealed associations between electrophysiological features and specific neuronal subpopulations, providing insights into the functional organization of the dorsal horn (DH). Finally, we demonstrate for the first time that the contralateral spinal cord encodes sensory information from the opposite body part in awake mice. Together, these findings provide insights into the functional organization and dynamics of the dorsal spinal cord, opening new perspectives for understanding spinal sensory processing.

## Results

### The Spinotrode: A novel mini-vertebral prosthetic implant for single-unit recording in freely moving mice

The spinal cord and surrounding vertebrae undergo mechanical stress during locomotion and natural behaviours^21^. Therefore, achieving stable, long-term, spinal recordings in freely moving animals without inducing tissue damage remains challenging. To address this limitation, we developed a novel implant that allows bilateral recordings from the deep layers of DH. Insulated nichrome microwires, combining suitable electrical properties, biocompatibility and small dimensions^34,35^, were used to minimize tissue impact and ensure reliability for long-lasting recordings (**Fig. 1a, left**). Two custom-made bundles incorporating eight insulated electrodes were embedded within 125-µm-thick guides of a biocompatible 3D-printed semi-vertebra. These paths are separated by 700 µm, allowing electrodes to be positioned at equivalent distances lateral to the central vein (**Fig. 1a, right**). To avoid an oversized device mounted on the back of the animal and to prevent the animal from chewing connectors, electrodes were connected to an Omnetics connector positioned on the skull with a biocompatible silicone sheath securing the wire bundles subcutaneously (**Fig. 1b-c**). This configuration stabilizes the electrodes within the spinal cord even as the spinal cord slides rostro-caudally relative to the vertebrae and implant (**Fig. 1c-e**). Finally, electrode tips were cut at a 45° angle, enabling recordings at different depths between 200 and 250 µm, reaching the deep layers of the DH (laminae IV-V) (**Fig. 1b, f**).

**Figure 1:**
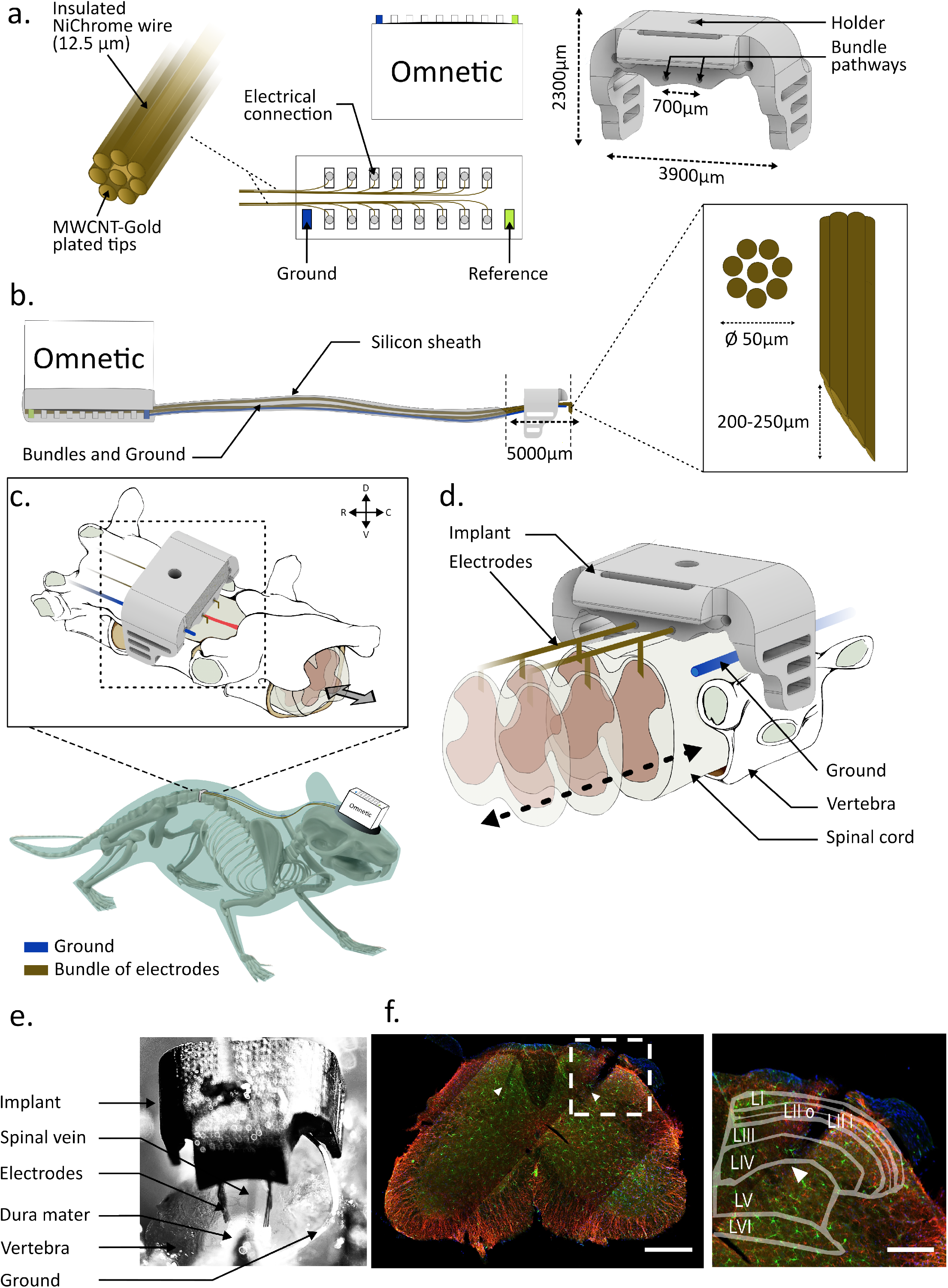
A novel implant for investigating spinal cord activity in freely moving conditions. a. Principal components of the novel implant. Left: Eight insulated Nichrome wires (external diameter: 12.5 µm) are bundled together to form an 8-channel recording electrode; two such recording electrodes are assembled. An 18-pin Omnetics connector is used as a miniaturized electrical interface between the implanted electrodes and the external recording device. Right: 3D-printed artificial vertebral implant designed to securely position electrodes above the spinal cord. Each 8-channel electrode slides freely through a pathway (external diameter: 125 µm) so that the implant, fixed to the vertebrae, can move rostrocaudally relative to the spinal cord without disrupting electrode placement. b. Graphical representation of the complete implant: the ground electrode and the two 8-channel electrodes are covered with a biocompatible silicon sheath. The allowed rostro-caudal movement of the 3D-printed component is indicated by the black arrow (∼5000 µm). Insert: schematic view of the tips of each electrode bundle (external diameter: ∼50 µm) cut at a 45° angle. c. Schematic representation of the implant location after surgery: the 3D-printed component is placed on top of a vertebra following a hemi-laminectomy. Bundles of electrodes (gold) are implanted bilaterally in the L5 dorsal horn of the spinal cord. The ground electrode (blue) is positioned near the tissue between L4 and L5 vertebrae. Wires are routed beneath the back skin, and the Omnetics connector is secured to the skull with dental cement. d. Zoomed view showing spinal cord movement relative to the vertebra with the electrodes implanted in the dorsal horn, facilitated by the 3D-printed component secured on top of the vertebra. e. Image of the implantation of two 8-electrode bundles carried in the 3D-printed component. f. Left: histological image of the lumbar dorsal horn with the two electrodes traces. Arrowheads show the tip of each bundle. (scale bar: 200 µm), Right: zoom-in of one of the two bundle trace (scale bar: 100 µm).

Overall, our implant addressed the challenges described above, enabling bilateral spinal recordings without affecting the animal’s well-being, locomotion (ANOVA F(5.0, 88.0) = 1.236, p = 0.2993) (**Fig. S1a)** or sensitivity to mechanical (2way ANOVA F(4.0, 16.0) = 1.271, p = 0.4443) and thermal (ANOVA F(1.472, 5.005) = 2.258, p = 0.1985) stimuli (**Figs. S1b, S1c, S2**). The small size of the electrodes minimized neuroinflammatory responses in the DH of the spinal cord (2-way ANOVA F(1.792, 4.301) = 2.2832, p = 0.7452) (**Fig. S1d, e**), making our approach a suitable technique for chronic spinal recording in freely moving mice.

**Figure 2:**
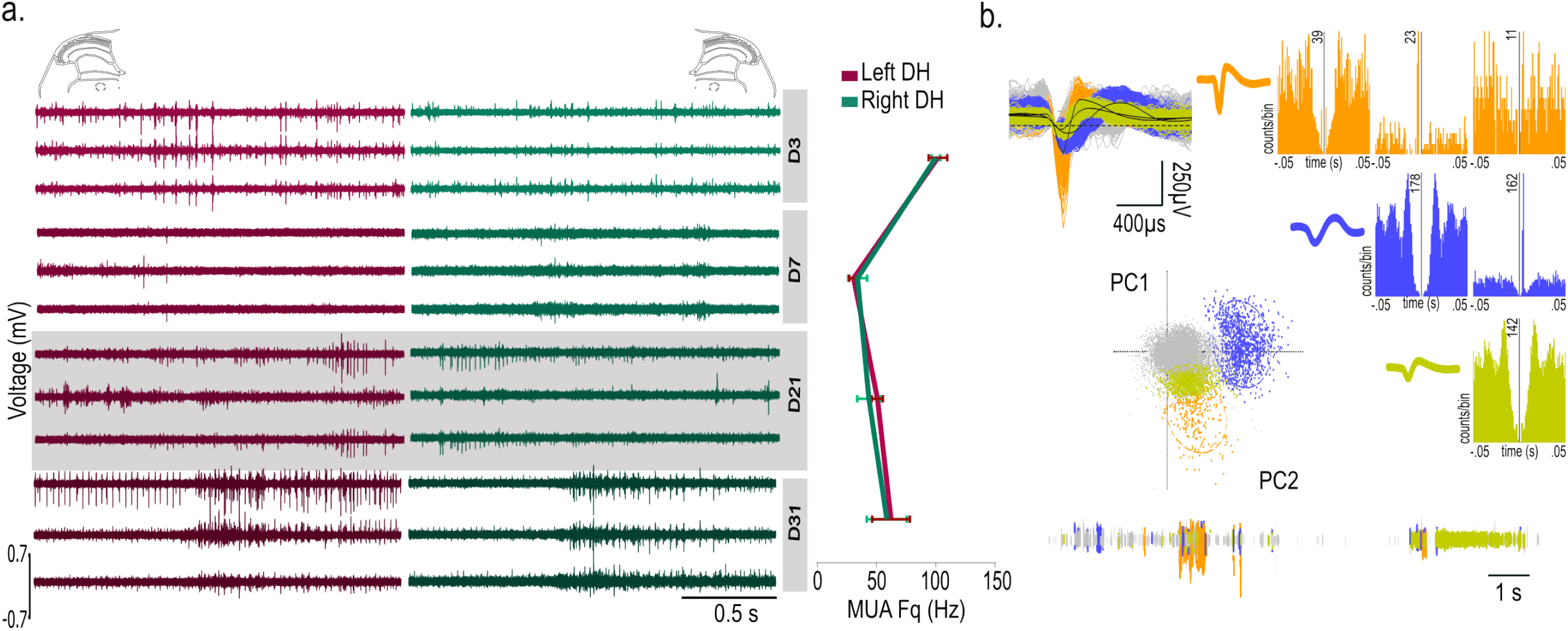
Bilateral long lasting and stable dorsal horn recordings in freely moving mice. a. Left: Sample raw signals recorded in the left (pink) and right (green) dorsal horns at 3, 7, 21, and 31 days post-implantation (3 channels/day). Right: multi-unit activity (average ± SEM) among the recording sites (up to 8) during 200 s in a freely moving animal, across days. b. Representation of the sorting method of putative DH units. left: Principal component analysis (PCA): associated waveforms (top),PCA-based clustering (middle), and the sorted waveforms from a single channel in the DH. right: auto-correlograms and cross-correlograms of sorted units and their associated waveforms.

### Achieving bilateral, long-lasting and stable DH recordings with the Spinotrode

Following successful implantations, we recorded spinal activity bilaterally from all available channels, with an average of 16 neurons in the left DH and 15 neurons in the right DH, respectively (**Fig. S3a**). Weekly recordings from day 3 to day 31 post-surgery showed high signal stability in both DH (**Fig. 2a**). Although, the mean impedance of all active channels increased post-implantation (ANOVA F(2.108, 156.0) = 50, p<0.0001), it remained stable across both sides of the spinal cord over weeks (F(3.0, 30.0) = 0.76, p = 0.52), maintaining high-quality recordings throughout (**Fig. S3b, c**). Interestingly multiunit activity peaked at day 3 (102.25Hz left, 100.50Hz right), decreased one week after implantation (30.7Hz left, 34.09Hz right) before increasing again at day 21 (50.92Hz left, 43.78Hz right) and day 31 (62.53Hz left, 58.90Hz right), in both DHs without significant difference (F (1.0, 52.0) = 0.7 568, p = 0.3883) (**Fig. 2a**). This signal stability across weeks allowed unit sorting in all suitable channels (see Methods). Distinct waveforms were then segregated in all mice using Principal Component Analysis (PCA), a standard method for waveform classification^36–38^, and validated by their auto-correlograms and waveform features (**Fig. 2b**).

Together, these results demonstrate that our device allows reliable recordings of a substantial proportion of DH units bilaterally in all implanted mice, thereby enabling in depth, long-term single cell analysis of the recorded signals.

### Bridging anesthetized and freely moving neural data by recording DH activity across motion states

It is well established that the spinal cord can be functionally segmented, with the DH processing sensory inputs and the ventral horn driving motor outputs according to *ex vivo* studies and experiments in anesthetized animals^39–41^. Our implant allowed us to track DH units across anesthetized, awake but immobile, and freely moving states and to compare raw and single-unit spinal activity between these conditions (**Fig. 3a**). Although some neurons are spontaneously active under anaesthesia, the overall population remained largely silent (0.53Hz ± 0.18). Interestingly, in awake, immobile mice, the overall population firing rate remained unchanged (0.75Hz ± 0.18) but rose significantly during locomotion (1.50Hz ± 0.32) (**Fig. 3b**). Yet, neuronal activity varied heterogeneously as a function of animal movement (**Fig. 3c**): 52% of recorded neurons were activated during movement (speed > 0.2m/s), 8% were inhibited, and 40% showed no difference in activity between immobility and locomotion (**Figs. 3c-d**). Among the activated neurons, 26% exhibited a positive correlation with speed (Pearson’s r > 0.15), indicating movement-related encoding in the DH of the spinal cord (**Figs. 3c-d**). These results show the state-dependent dynamics of DH neurons with some units, albeit a minority, increasing their activity during locomotion.

**Figure 3:**
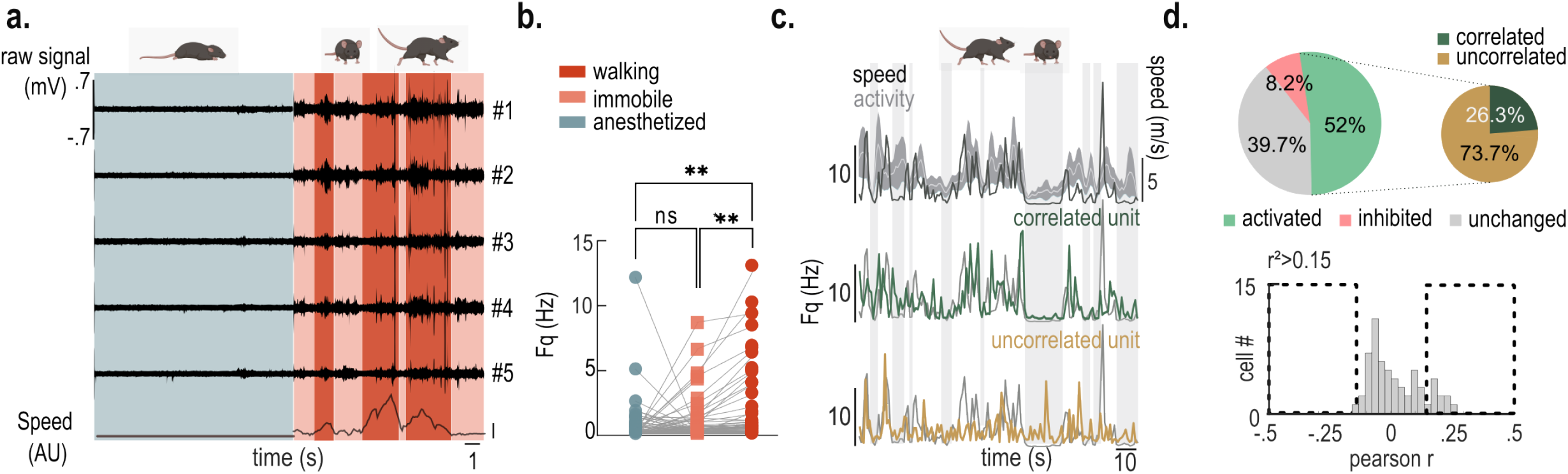
Dorsal horn neuronal activity reflects the animal’s motion state. a. Raw traces from five recording channels in a single animal while anesthetized (gray), awake but immobile (orange), or walking (red). The bottom trace represents the animal’s locomotor speed. b. Single unit firing frequency (Hz) of the overall population (3 mice, 73 units) during anesthesia, immobility, or walking (ANOVA F (3,72) = 16.71, p = 0.0002, post hoc Dunn’s paired comparison, significance *p < 0.05 **p < 0.01, ***p < 0.005). c. Top: Graphs of average firing frequency (Hz, mean ± SEM, n = 15 units) superimposed with the animal’s speed (m/s); middle: example of a unit whose activity correlates with speed; bottom: example of an uncorrelated unit. d. Top: pie chart showing the distribution of neuronal populations that are activated (52%), inhibited (8.2%) or unresponsive (39.7%); inset pie chart showing the percentage of cells positively correlated or uncorrelated cells with speed among the activated cells. Bottom: histogram showing Pearson correlations between the firing activity of individual units and the animal’s speed (3 mice, 74 units). Positively correlated units, r > 0.15, p < 0.05 and negatively correlated units r < −0.15, p < 0.05) are indicated

### Revealing relationships between sensory input, DH neuron responses, and withdrawal behaviour

A major advantage of the Spinotrode to decode nociception is the ability to relate DH neuron activity to nociceptive behaviour. We thus quantified the response of DH neurons to mechanical and thermal stimulation of the paw in awake animals while monitoring their nociceptive withdrawal reflex. To achieve this, we used Spinotrode recording coupled to a time-sensitive, semi-automated robotic system, RAMalgo^42^, to deliver controlled mechanical and thermal stimuli ranging from innocuous to noxious intensities while animal behaviour was videotaped, and paw withdrawal time was precisely measured (see Methods)^43^. This approach enabled correlation of DH activity with ipsilateral hind paw stimulation (thermal or mechanical) from onset to paw withdrawal (PW) and beyond (**Fig. 4a, b).** We plotted the input-output relationship between the firing rate of each identified unit and the stimulus from onset to time of PW (**Fig. 4c, d).** Of 154 recorded units, 70 were activated by mechanical stimulation and 51 by thermal stimulation of the hind paw. No units were inhibited by stimulation. A proportion of these units responded to both thermal and mechanical stimulation (37 units; **Fig. 4e**). We identified two subsets of cells based on their response dynamics to the stimulus intensity. Units with a low threshold and no correlation with input intensity were classified as non-linearly responding (NL) units^41^ (**Fig 4f-k**). A second class of neurons responded with graded levels of activity throughout the range of stimulus intensity and were labelled as linearly responsive (LR) units, with input-output profiles corresponding to that of wide dynamic range (WDR) neurons when responding to controlled graded intensity input^41^ (**Fig 4f-k)**. The LR pool of cells also displayed graded linear responses at the population level (**Fig 4j-k**, bottom panel). PW events occurred at variable latencies after paw stimulation, with latencies of 0.2-6s for mechanical stimuli and 0.1-20s for thermal stimuli (**Fig. 4l-m)**. To precisely observe neuronal response at the onset of PW, we aligned each neuronal response to the PW. At the level of the population, we observed recruitment of units leading up to PW, a peak at PW, and a proportion of units discharging after PW (**Fig. 4n**).

**Figure 4:**
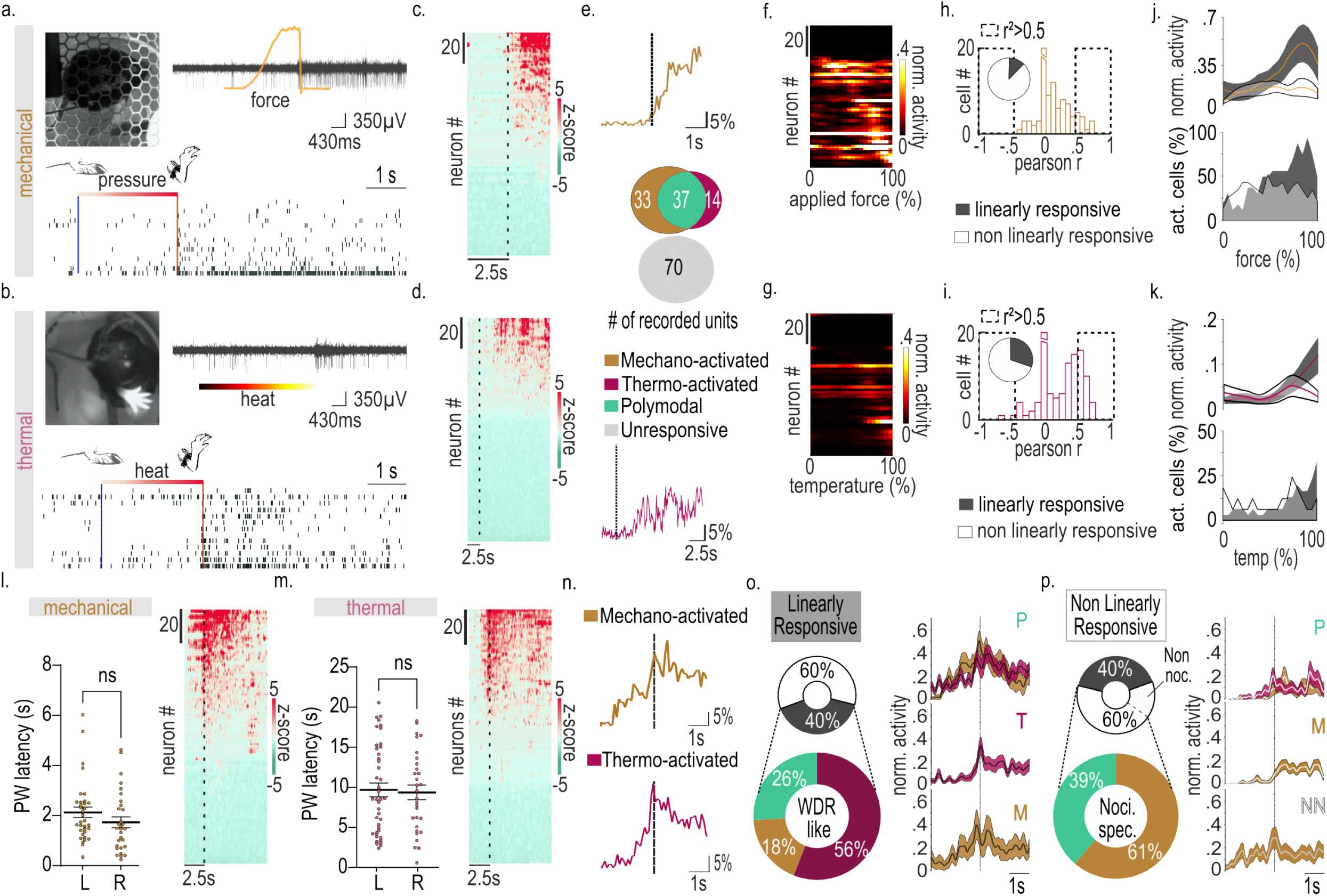
Dorsal horn neuronal response to mechanical and thermal stimuli in freely moving mice. a.,b. Top Left: Video frame during a mechanical or thermal trial using RAMalgo^42^. Right: raw trace of a single-channel DH activity, aligned with the force trace or temperature increase until paw withdrawal. Bottom: raster plot of unit responses from a single mouse (13 cells) to mechanical or thermal stimulation of ipsilateral paw; blue line shows stimulus onset and red line indicates paw withdrawal. c.,d. Heatmaps of the overall cell population activity (z-scored, bin= 0.05 s, n=154, 6 mice) during mechanical (c) or thermal stimulation (d). Ipsilateral neuronal responses are aligned to stimulation onset and units are sorted decreasingly according to their z-score activity. e. Time course shows the percentage of cells activated per bin (0.1 s) in response to mechanical (top)(−2.5 to 2.5 s) and thermal (bottom)(−2.5 to 10 s) stimuli among recorded cells. Euler Venn diagram shows distribution of neurons based on their sensory profile (n = 154, mice = 6). Values are expressed as the number of recorded cells per category. f.g., heatmaps of sequential normalized activity for each sensory neuron to increased applied force (f) and temperature (g). h.,i. Histograms of Pearson correlation coefficient between neural activity and force (h) or heat increase (i). Insets show the proportion of linearly and non-linearly responsive units, pressure: 12% (h), heat: 30.5% (i). j.k., Graphs combine normalized activity (average ± SEM, from 0 to 1) (top) and the percentage of activated cells (bin step = 0.10s for pressure and 0.05s for heat) (bottom) of the linearly responsive (grey) and non-linearly responsive cells (white). Activity is correlated to increased applied force and temperature before withdrawal (100%). l.m., left: all withdrawal latencies (mean ± SEM) for all animals (n=6) for the left and right paw during mechanical (l) and thermal stimulation (m) Right: heatmaps of the overall cell population activity (z-scored, bin= 0.05 s, n = 154, 6 mice) during mechanical (l) or thermal stimulation (m). Ipsilateral neuronal responses are aligned to paw withdrawal (PW) and units are sorted decreasingly according to their z-score activity. n. proportion of activated cells per 0.1s time bin around paw withdrawal during ipsilateral mechanical (top) or thermal (bottom) stimulation. o.p. left: Sunburst pie chart illustrating the distribution of sensory response profiles amongst linearly responsive neurons (o.,grey), which are all wide dynamic range, and amongst non-linearly responsive cells (p), which include nociceptive specific (84%) and non-nociceptive (16%). Right: average normalized activity ± SEM response to mechanical (yellow) or thermal (purple) stimulation, aligned to paw withdrawal (−2.5 to 2.5s) and subdivided by cell category (P: polymodals, T: thermo-activated, M: mechano-activated, NN: non-nociceptive).

We identified three response patterns: one population comprising LR units started responding before PW, reached their peak response at PW, and continued firing after PW, and were labelled WDR-LR; another population comprising NL units with peak activity at PW were deemed nociceptive specific and labelled NL-NS; a separate population of NL units with variable peaks of activity before PW were deemed non-nociceptive and labelled NL-NN (**Fig. 4l-p and S4**). We separated the WDR-LR, the NL-NS and the NL-NN, to study their response at PW. Among the population of cells active beyond PW, 40% were WDR-LR (56 % thermal-only, 18% mechanical-only, and 26% polymodal; **Fig. 4o**) and 60% were NL (50.4% NL-NS and 9.6% NL-NN. For NL-NS: 61% mechanical-only, 39% polymodal; **Fig. 4p**).

We sought to sort neurons on the basis of their discharge frequency and waveform characteristics, namely spike half-width (SHW) and area under the curve (AUC). We identified three clusters that differed in AUC and SHW: SHW increased progressively from cluster 1 to cluster 3 (197, 340.9, and 529.5µs), cluster 2 displayed the largest AUC (135.46 mV^2^), whereas mean frequency was similar across clusters (**Fig. 5a-b**). We compared their response profiles at PW (thermal and mechanical) to the different functional categories (z-score **Fig. 5c and Fig. S4**). Clusters 1 and 3 units were not associated with any specific type of stimulus (mechanical or thermal) or response characteristics (LR vs NLR). By contrast, cluster 2 was highly enriched in NL-NS mechanical units (**Fig. 5d-f**).

**Figure 5:**
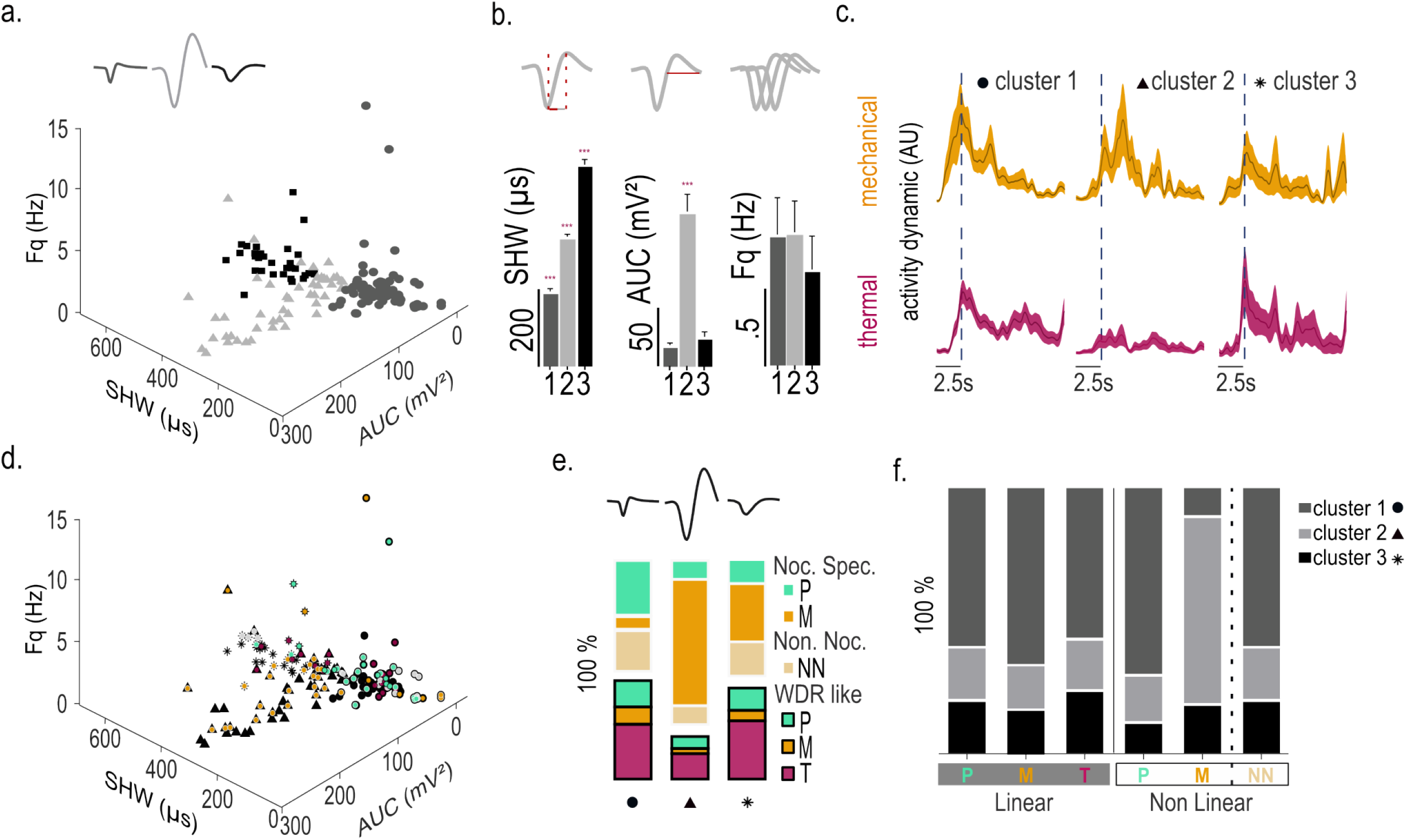
Functional mapping of dorsal horn neurons based on their waveform properties and sensory profiles. a. 3D scatter plot of waveform-based clustering showing three clusters identified using k-means: cluster 1 (squares, n=46), cluster 2 (triangles, n=32) and cluster 3 (circles, n=18). b. Bar graphs showing the mean ± SEM of spike half-width (µs) (ANOVA F(1,826, 73,96) = 330.7, p < 0.0001), area under the curve (mV²) (ANOVA F (1,084, 43,89) = 58,02, p < 0.0001), and firing frequency (Hz) (ANOVA F(1,864, 140,8) = 0,2463, p = 0.7666) for each cluster. Bonferonni post hoc test * different from all groups, *** p < 0.001.c. Activity patterns (z-score average ± SEM, bin = 0.05 s) for each cluster aligned to paw withdrawal (−2.5 to 10 s) during mechanical (top) and thermal stimulation (bottom); cluster1 (n=46), cluster 2 (n=32), cluster 3 (n=18). d. 3D scatter plot of the three clusters (symbols as above) overlaid with colors indicating sensory profile. e. Distribution of sensory profiles across clusters. f. Distribution of clusters across sensory profiles. (P: Polymodal, M: Mechanical, T: thermal, NN: Non Nociceptive).

The results confirm previous characterization of linearly (WDR) vs non-linearly (NS) encoding neurons as a function of stimulus intensity (lavertu et al, Brain 2014). What the new approach reveals, for the first time, is that the activation of NL-NS effectively coincides with withdrawal reflex, demonstrating their key and specific role in nociception.

### Spinotrode parallel recording capability reveals DH bilateral encoding of a unilateral stimulus

We took advantage of our probe that allows bilateral spinal recordings to evaluate the contralateral neuronal response to mechanical and thermal stimulation. Unexpectedly, in awake behaving animals, we found that 57.8% of neurons responded to noxious mechanical stimulation of the contralateral paw, and 29.8% to a noxious thermal stimulation contralaterally (**Fig. 6a-d)**. This was never observed in anaesthetized animals (**Fig. S5**). In contrast to responses of ipsilateral DH neurons, responses to contralateral stimulation were associated with the onset of PW for both mechanical and thermal stimulation (mechanical: diff = −0,8577, t(10) = 5.915, p<0,0001; thermal: diff=-0,5958, t(10) = 4.968, p=0.0006) (**Fig. 6c-f**). Comparing the response profiles of the same recorded units to both ipsilateral and contralateral stimulations, we found that neurons that were LR to ipsilateral stimulation were largely NL to contralateral stimulation (**Fig. 6g**). The latter difference in response profile suggests that the contralateral response may be more related to the withdrawal than the sensory response. It is unlikely that the response was due to sensory input on the contralateral side due to the withdrawal event, because it was heterogenous in terms of modality and associated with nociception. (**Fig. S5**). This is further strengthened by the observation that some cells were activated only by contralateral stimuli.

**Figure 6:**
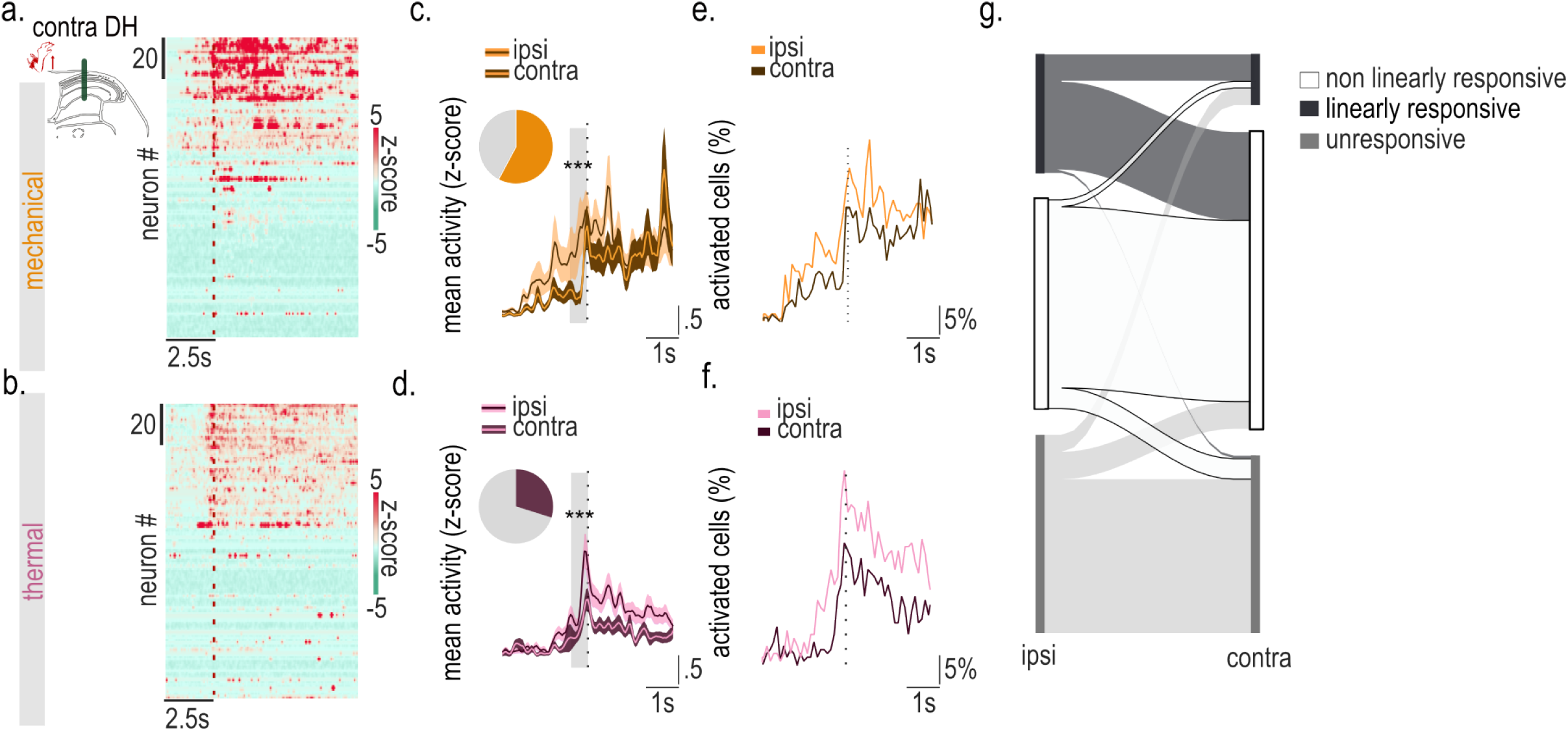
Bilateral encoding of unilateral sensory stimuli. a., b. Left: schematic of dorsal horn recordings during ipsilateral paw withdrawal (PW); Right: heatmaps of the overall cell population activity in the contralateral dorsal horn aligned to PW and ordered by decreasing z-score (z-scored, mean bin= 0.05 s, n=154, 6 mice) during mechanical (a) and thermal stimulation (b). c., d., Overall neuronal activity in the ispilateral dorsal horn (z-score ± SEM, bin = 0.05 s) overlayed with the contralateral side, aligned to PW (−2.5 to 2.5 s) during mechanical (c) and thermal stimulation (d). e.,f.,Proportion of activated cells in both dorsal horns (bin = 0.1 s), aligned to PW (−2.5 to 2.5 s) during mechanical (e) and thermal stimulation (f). g. Sankey plot illustrating the neuronal dynamics across both dorsal horns (n = 154) for linearly and non-linearly responsive cells in response to an ipsilateral and contralateral paw stimulation (Chi²: df= 14.14, 2, p<0.0009).

Our results reveal, for the first time, functional responses in the sensory DH to contralateral nociceptive input. Because it was preferentially associated with the withdrawal response, it suggests the existence of widespread complex nociceptive coding. Importantly, this was only observable in unanaesthetised animals, yet could not simply be associated with movement given the functional types of cells activated, including a large proportion of NS cells.

## Discussion

We introduced an innovative device for longitudinal recordings of spinal DH activity in freely moving animals. By minimizing tissue shear and backload, the Spinotrode preserves physiological posture and naturalistic motion while enabling stable recordings over several weeks. The Spinotrode thus enables the neural basis for sensorimotor function to be interrogated with single-cell resolution.

To date, few studies have developed tools capable of reliably accessing such spinal activity in behaving animals. Early studies using microelectrodes in rats and cats revealed differences between anesthetized and awake animals and neuronal plasticity during goal-directed behaviour but were restricted to single experiments^26^ and did not allow natural locomotion^3,32^. More recently, new in vivo approaches enabling recordings in awake and freely moving rodents have emerged^22,24,25,30,31,34^, but most of these have been limited to recordings from superficial laminae at only one site. Our device overcomes these limitations by enabling simultaneous bilateral recordings from multiple neuronal populations at different depths, while preserving natural movements and ensuring long-term stability. Additionally, the strength of this novel device lies in its scalability to recordings from multiple segments by adding spinal vertebral implants. The introduction of a customizable 3D-printed vertebral implant also opens avenues to also incorporate microLEDs for epidural optogenetics^35,36^.

The Spinotrode allowed us to reveal that spontaneous DH activity is modulated by animal movements, but is mostly uncorrelated with locomotor speed. Yet, we still identified a subset of units closely correlated with speed of movement, suggesting premotor functions in the DH traditionally considered purely sensory^37,38^.

A key finding enabled by our Spinotrode recordings is the relationship of neural activity with reflexive withdrawal, the key indicator of a nociceptive response. The results confirmed previous classification of neuronal types (NN, WDR, and NS), but in particular, confirms the key association between nociceptive specific (NS) neurons and the nociceptive reflex. Both WDR and NS cell types also display a continuous after-discharge following nociceptive withdrawal, implicating those cells in complex nociceptive behaviours following withdrawal. Finally, bilateral recordings revealed contralateral activation of a significant subset of cells, specifically upon the nociceptive withdrawal event and beyond, uncovering previously ignored role for DH nociceptive circuit in coordinating the after response to complex nociceptive behaviours. The latter sensory activity is unlikely to be a simple response to movement as it is specifically related to the nociceptive regime with no correlation to movement.

As the Spinotrode is easily modifiable, increasing the electrode length and the spacing between recording sites offers the possibility for simultaneous recordings in the dorsal and ventral horns, thus enabling analysis of sensory-motor integration. Our results also allowed us to associate waveform properties to functionally classified subpopulations, namely, narrow spikes (i.e. short SHW) were predominantly associated with polymodal neurons, and strongly biphasic spikes (i.e. large AUC) were associated with mechano-nociceptive neurons. Relationship between the waveform and neuron types has been amply documented in brain regions but also in the DH of the spinal cord^44,45^ (e.g., excitatory and inhibitory neuron spikes segregated by their monophasic vs. biphasic waveforms, respectively DH^45^). Our method thus opens a path for fine-scale microcircuitry analysis and the dynamics of gating and disinhibition in the spinal cord, in freely moving animals^46^.

Lastly, current understanding of spinal mechanisms is largely focused on ipsilateral DH responses to sensory inputs^47,48^. However, in behaving animals, proprioceptive and motor signals are closely associated with somatosensory and nociceptive inputs, reflecting integrated lateralized processing. Thanks to our Spinotrode, we were able to analyse the neuronal response evoked by contralateral mechanical and thermal paw stimulation in awake behaving conditions. Our results revealed a similar response to the ipsilateral side with some units, mainly thermal, modifying their response to the nature of the peripheral stimulations of the paw. Such findings force a reassessment of sensory motor interactions, and especially to consider the implication of the DH in the control and coordination of complex nociceptive withdrawal responses.

The novel approach we introduced has the potential to revolutionize spinal electrophysiological studies. By enabling high-resolution monitoring in awake animals, it uncovers unprecedented insights into spinal DH physiology, including the identification of premotor, sensory-specific, and polymodal units, as well as their dynamic responses to bilateral peripheral nociceptive inputs. Beyond its immediate applications, this technology opens new avenues for studying neural plasticity in the spinal DH following peripheral or central lesions, linking cellular activity and behaviour, a key step towards identification of the cellular and circuit underpinnings of somatosensory processing in health and disease.

## Methods

### Animals

All experiments in this study were conducted in accordance with the regulations of the Canadian Council on Animal Care and the European guidelines. They were approved by the Ethics committees of the University of Bordeaux (APAFIS #32137) and University Laval (#2022-1038-3). Male C57BL/6J (#000664; JAX) aged between 6 and 8 weeks were housed in groups of 2-5 in collective cages on ventilated racks prior to implantation, and individually afterward to minimize loss or damage of implants. Animals had *ad libitum* access to food and water and were maintained under a standard light/dark cycle (7:00/19:00), at constant temperature (21 ± 2 °C) and humidity levels (60%). A one-week acclimation period was respected prior to implantation, and the animals were subjected to sensory testing only after a two-week recovery period.

### Implant fabrication

To suit the anatomy of the spine, the dimensions of a vertebra from a 10-week-old mouse were measured and used as a template for implant design. The vertebral implant was modeled using computer-aided design (CAD) software (Inventor Pro 2023, Autodesk) and its key features are shown in Fig.1a. To minimize tissue rejection and inflammatory responses, the vertebral implant was engineered using a microfabrication 3D printer with a 1.9 µm resolution (Fabrica Printer, Nano Dimension) and a biocompatible resin (Medical M-810, Nano Dimension) providing suitable robustness for long-term implantation. For electrode fabrication, ultrathin (12 µm external diameter) polyimide coated Nickel-Chrome (NiCr) wires (#RO800, Alleima) were selected to minimize biological impact. Wires of 15 cm length were bundled together using mineral oil (#M5904, Sigma-Aldrich) to assemble two independent 8-channel recording electrodes. Following bundling, each wire was individually bonded to a pin of an Omnetics connector (#A79042-001, Omnetics Connector Corporation) using water-based conductive silver paint (#842WB, Digikey). The distal end of each bundle was then inserted into the holes of the implant specifically designed to allow electrode movements, as illustrated in Fig.1b-d. A PFA-coated silver wire with a diameter of 76.2 µm (#785500, A-M systems) was used as the ground wire. Electrical insulation was then achieved by applying silicone (#3140RTV, Dow) to the connectors and along the bundles. Finally, the distal tip of the two bundles was trimmed at a 45° angle.

### Impedance tuning

Gold electrodeposition was performed to lower the impedance of each electrode. The electrodeposition solution was prepared following the protocol described by ^49^ by mixing 75% multi-walled carbon nanotube (MWCNT) solution (1 mg/mL in distilled water; 8 nm, Cheaptubes) with 25% cyanide-free gold solution (#31-0611-0100, Neuralynx). To facilitate MWCNTs dispersion, the mixture was sonicated prior to the addition of the gold solution. Electrodeposition was then performed using the NanoZ system and software (Neuralynx). The protocol from ^50^. was adapted to allow adjustment of electrode impedance to approximately 150 kΩ. An initial current of +0.100 µA was applied across all electrodes, followed by −0.100 µA for 3 seconds until the target impedance was reached. Impedance was gradually reduced in three steps: 500, 300, and 150 kΩ, with measurements taken at 1 kHz between each deposition step.

### Electrodes implantation

Eight mice were deeply anesthetized with an isoflurane-oxygen mixture (4% induction, 1.5-2% maintenance) and placed in a stereotaxic frame (#68025, RWD) on a heating pad, and the head was immobilized using ear bars. Mice received ocular gel, a subcutaneous injection of buprenorphine (0.5-1mg/kg), and a local injection of lidocaine-bupivacaine (volume max: 0.08mL/10g) prior to skin disinfection.

The skull was exposed, and a 1-2 cm incision was made in the dorsal skin to expose the lumbar spinal region. The implant was carefully inserted through the incision, and a micro-hole was then drilled for the insertion of a microscrew (#19010-11, FST) to improve connector fixation on the skull. The connector was then secured using dental cement (#186-1068, C&B Metabond).

After identification of the T12-L2 vertebrae, a small incision was made between the tendons to expose these vertebrae prior to fixation of T12 and L2 using spinal clamps (#505289, WPI). The spinous process of the L1 vertebra was removed, and the dorsal surface was flattened. A hemilaminectomy was then performed on its caudal portion. Finally, the dura mater was incised bilaterally using a 30G needle to facilitate insertion of the electrode bundles. Electrodes were bilaterally inserted into the spinal tissue along the dorso-ventral axis in 10µm increments using a stereotaxic micromanipulator to minimize tissue damage. Spinal electrophysiological activity was continuously recorded using the OpenEphys system (#OEPS-9029, OpenEphys) to guide placement. To confirm correct positioning, slight pressure was applied to the hind paws during electrode insertion. Once stable bilateral evoked responses were observed, the implant was secured to the vertebra with dental cement, avoiding contact with the spinal cord and the skin was sutured.

### Assessment of nociceptive sensitivity

Six mice were used for the sensory behavioural experiments following surgery. All behaviour experiments were conducted between 08:00 and 14:00. To assess mechanical sensitivity, animals were placed in a small cage with a mesh grid on the floor and were allowed a 30-minute acclimation period prior to testing. The Simplified Up-Down (SUDO) method was used as described by Bonin et al ^51^., with von Frey filaments #2 to #10. Paw withdrawal thresholds (PWTs) were converted to force (g) using the following equation:

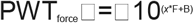

where F is the filament number, and *x* and B are constants depending on the filament range. For filaments #2 to 7, *x* = 0.240 and B = −2.00; for filaments #7 to 10, *x* = 0.182 and B = - 1.47.

Thermal sensitivity was assessed using the Hargreaves test as described by Hargreaves et al.^52^. Briefly, a calibrated infrared heat stimulus (IITC Life Science) was applied to the plantar area of the paw until withdrawal. Withdrawal latency was measured in seconds. The infrared intensity was set to 25, and a cutoff time of 25 seconds to prevent tissue injury. Five trials were performed on each paw, and the mean value was calculated. Mechanical and thermal sensitivity were assessed one week prior to implantation and weekly after surgery.

### Iba1 immunohistochemistry and quantification

Mice were euthanized and perfused through the left ventricle with 4% paraformaldehyde (pH 7.4). L1-S1 spinal cord segments were selected, post-fixed in the same solution and cryoprotected overnight at 4°C in 30% sucrose prepared in 0.1 M PBS. Coronal sections (30 µm) were cut, and free-floating sections were permeabilized in 0.1 M PBS containing Triton X-100 (0.3%) and bovine serum albumin (1%, Sigma-Aldrich) for 30 minutes. After washing, slices were incubated overnight with rabbit anti-Iba1 antibody (1:2000, #019-19741, Wako). The following day, immunoreactivity was revealed using an anti-rabbit peroxidase EnVision system (DAKO) followed by DAB incubation. Sections were mounted, cover-slipped, dehydrated and scanned at x20 magnification using a high-resolution scanner (Panoramic Scan II, 3DHISTECH Ltd, France). Images were analysed using Fiji/ImageJ software using custom scripts. Briefly, after background subtraction, a detection threshold was manually set to isolate Iba1 labelling, and a binary mask was generated from this signal. Microglial surface area was then quantified in both DHs of the spinal cord, which was manually defined.

### RAMalgo (reproducible automated multimodal algometer)

To correlate spinal neuronal activity with behavioural responses, we used the RAMalgo system developed by Dedek et al.^42^. This system enabled the delivery of thermal and mechanical stimuli to the plantar surface of the hind paws with high temporal synchronization with electrophysiological recordings. Aiming is conducted via computer using video and motorized actuators, thus allowing the human experimenter to remain away from the tested animal. All trials are recorded on video synchronized to stimulation and other measurements (see below), thus enabling post hoc validation of stimulus onset and paw withdrawal timing.

Mechanical stimuli were delivered using a dual-mode articulated arm (300C-I, Aurora Scientific), allowing precise control of probe displacement, velocity and force. Stimulation parameters were set manually via the interface. Force applied on the hind paw was continuously monitored at 1 kHz and paw withdrawal was automatically detected as a rapid drop in applied force, triggering immediate termination of the stimulus. Behavioural responses were recorded for an additional 20sec following withdrawal to correlate spinal neuronal activity with delayed nociceptive behaviours.

Thermal stimulation was delivered using infrared laser (980nm, Laserglow Technologies). Laser intensity was tunable via the interface, with multiple levels tested (1.25 - 2.50V), ranging from innocuous to withdrawal-evoking stimuli. Maximum stimulation duration was limited to 17 seconds to prevent tissue damage. Paw withdrawal was automatically detected using a red LED (625 nm) coupled to the laser. Red light reflected from the paw was measured in real time by a photodetector positioned beneath the animal and sampled at 1 kHz. A user-defined detection threshold was set at 90%; a decrease in reflected light exceeding 10% triggered simultaneous termination of the laser and LED, marking paw withdrawal.

### In vivo recordings in freely moving animals

#### Data acquisition

To connect the headstage to the Omnetics connector, mice were lightly anesthetized with an isoflurane-oxygen mixture (4%). Neuronal activity was recorded using a 16-channel amplifier headstage (#C3334, Intan Technologies) with ADC sampling at up to 30 kSamples/s per channel. Signals from the probe were acquired using the OpenEphys recording system (#OEPS-9029, OpenEphys). Before recording, channel impedance was measured, and channels with values above 2 MΩ were excluded. Signals were processed through a signal chain, bandpass filtered (300 - 6000 Hz), and referenced to the common average, with the signal averaged across the eight electrodes of each bundle. To ensure synchronization between electrophysiological recordings and stimulation, an input/output board (#OEPS-6501, OpenEphys) was used to connect the acquisition unit to the 1401 acquisition interface (Cambridge Electronic Design), which controls the RAMalgo stimulation system.

Single unit segregation was performed using Plexon Offline Sorter software. All behavioural recordings were performed the same day without unplugging the mouse, so the files were concatenated in a single file in order to perform unit sorting. Waveform clusters were manually defined using principal components, time and voltage features of the waveforms. Clusters were defined as isolated from the background noise in the feature space with no spike within a refractory period less than 1.5 ms, after checking the auto-correlograms. Cross-correlograms were also performed, any unit displaying a peak of co-activity at 0ms on other channels was considered as a duplicate. The duplicate with the higher number of spikes was removed.

### Data Analysis

#### Waveform property classification

The defined DH single units were classified based on similarities in their waveform shape using three criteria: spike half-width (µs), the area under the curve, and the firing frequency (Hz)^44^. The distance similarity of combined features to assign neurons to the closest cluster was computed with K-means. The optimal number of clusters (3 in this case) was defined using the silhouette function, iterated one thousand times for robust accuracy, and the value of k with the highest mean silhouette score was selected.

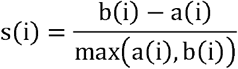

- a(*i*): mean distance between *i* and all points of its own cluster (cohesion)
- b(*i*): mean distance between *i* and all points from the closest cluster (separation)

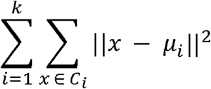

- k = number of clusters,
- C*_i_*= set of data points in cluster *i*,
- µ*_i_*= centroid (mean) of cluster *i*,
- ∥x−μi∥^²^= squared Euclidean distance between a data point *x* and its cluster centroid µ*_i_*

##### Longitudinal multi unit activity analysis

In order to compare the overall neuronal activity of the signal between several days, we compared the mean frequency of all suprathreshold spikes of the filtered signal, of all good channels (animal n=1), for 200s. We separated left and right DH channels and compared them at day 3, 7, 21, and 31. We performed a 2-way ANOVA with a Bonferroni post hoc test.

##### Heatmaps and single cell classification

Cell classification was assessed for each unit by defining its response significance to an event (heat, mechanical stimuli, and paw withdrawal). We calculated the average frequency of a baseline period 2s right before the event compared to a period at event onset (a 10s period from stimulation onset, and a 2s period before stimulation onset). To compare whether the activity was significantly lower or higher than baseline, we measured the difference of the two-sample using Cohen’s U3-test, with a p-value < 0.01, bin=0.050s.

The cell classification was used for grouping the z-score means by response type, for the creation of proportion graphs, Euler-Venn diagram and Sankey diagram.

Heatmaps and activity visualisation of correlating neural data around a specific event (pressure start, heat start, paw withdrawal) was calculated as the firing rate with a binned spike count of a 0.050s window. expressed as a z-score. To reduce bin-to-bin variability, the binned firing rate was smoothed using a two-dimensional Gaussian kernel (σ = 2) or a locally estimated scatterplot smoothing (LOESS, span = 0.03),

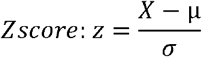

- X = individual data point,
- µ = mean firing rate over the 2 s baseline interval preceding stimulation
- σ = standard deviation of firing rate during baseline interval.

Alternatively, activity was normalized from 0 to 1 (see *“Normdata”* below).

##### Continuous cross-correlation to speed, heat and pressure

To evaluate the correlation between neural data and a continuous feature, we computed a Pearson’s correlation between firing rate and the stimulus temperature, stimulus force, or locomotor speed during corresponding intervals.

Speed: spike count interval and speed interval of 0.5s. The neurons were considered positively or negatively correlated with r²>0.15 and p-value<0.01.

Force and temperature: we normalized the concatenated trial traces from 0 to 1, with 0 as the minimum bin count value of the whole session (heat stimulation or mechanical stimulation comprising 2 to 6 repeated trials for each paw) and 1 as the maximum.

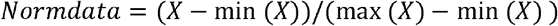

The neurons were considered positively or negatively correlated with r²>0.5 and p-value<0.01.

For pressure, if not correlated, neurons were separated by their mean firing activity being below or above 0.02 for nociceptive specific (NS) and non nociceptive (NN), respectively.

### Statistical analysis

For behavioural and histological analyses, comparisons between means, proportion graphs and histograms were performed using GraphPad Prism (version 10.4.1). Groups were compared using one-way or two-way ANOVAs followed by appropriate post hoc tests. Data are presented as mean ± SEM, and differences were considered statistically significant at p < 0.05. Neural data analysis was computed using MATLAB custom scripts (MathWorks, license # 41250949) the appropriate significance tests are presented in each *data analysis* subsection.

## Acknowledgements

We would like to thank Jonathan Lesveque from Benoit Gosselin’s team at Laval University and David Shrayber from Nano Dimension for kind help in 3D printing. We also thank the Bordeaux Imaging Center, BIC, UMS, CNRS, Université de Bordeaux and the Histocare platform, IMN CNRS UMR5293.

## Author Contribution

J.V., P.F., YdK conceived and designed the experiments; J.V., L.B., M.J., C.D., collected data; R.B.B. ethical authorization and immunohistochemistry. J.V., L.B., M.J., performed analysis; C.D., S.P., B.G., developed tool analysis; L.B., J.V., Y.dK., P.F., wrote the paper; B.G., S.P., A.B., F.W., carefully read and edited the manuscript.

