## Supplementary material for "Spinotrode: long-term intraspinal electrophysiological recordings to unravel dorsal horn neuron dynamics in behaving mice": Sup Figure Legends

**Supplementary Figures**

**Figure S1: Unilateral dorsal horn implantation preserves motor and sensory functions with no detectable post-surgical inflammation**. a. Distance travelled in an open field (cm) before the surgery and weekly after implantation (mice = 25 to 5), (ANOVA F(5, 88) = 1.236, p = 0.2993). b. Nociceptive withdrawal threshold to mechanical stimulation of the hind paw prior to implantation and weekly after the surgery (mice = 25 to 5), (2way ANOVA F(2.431, 8.751) = 1.038, p = 0.4082). c. Nociceptive withdrawal threshold to thermal stimulation of the hind paw prior to implantation and weekly after the surgery (mice = 25 to 5), (2way ANOVA F(1.472, 5.005) = 2.258, p = 0.1985). d. Illustrate image (left) and magnificance (right) of Iba1 immunostaining in the dorsal horn of the grey matter of the lumbar spinal cord. Arrowheads show labelled mibroglia (scale bars: 200 µm and 20 µm). e. Quantification of microglial surface and Iba1 ipsilateral-to-contralateral ratio in the dorsal horn of the spinal cord at different weeks post implantation (mice = 4 to 5), (ANOVA F(4, 18) = 2.901, p = 0.0514). All values are presented as mean ± SEM.

**Figure S2: Sensory function remains unaffected after bilateral dorsal horn implantation.** a. Nociceptive withdrawal threshold to mechanical stimulation of the hind paw prior to implantation and weekly after the surgery (mice = 5), (2way ANOVA F(4, 16) = 1.271, p = 0.4443). b. Nociceptive withdrawal threshold to thermal stimulation of the hind paw prior to implantation and weekly after the surgery (mice = 5), (2way ANOVA F(1.792, 4.301) = 2.2832, p = 0.7452). All values are presented as mean ± SEM.

**Figure S3: Dorsal horn recordings with stable impedance over weeks.** a. Number of neurons recorded per dorsal horn and per mice (n = 242, mice = 8), (T-test F(7,7) = 2.474, p = 0.2550), and their associated waveforms (bottom) b. Electrode impedance (mean ± SEM, MOhms) of each active channel before implantation and across weeks post-implantation in both dorsal horns (mice = 6), (ANOVA F(2.108, 156.0) = 49.99, p < 0.0001), Tukey's post hoc test * p < 0.05, **** p < 0.0001. c. Average electrode impedance (mean ± SEM, MOhms) across active channels per mice on both sides of the spinal cord, during implantation and weekly post-surgery (mice = 6), (2way-ANOVA F (3.30) = 0.7626, p = 0.5240), Šídák's post hoc test * p < 0.05, ** p < 0.01, *** p < 0.001. All values are presented as mean ± SEM.

**Figure S4: Dorsal horn neurons show linear and non linear activity pattern in relation to the increasing force applied.** a. Graphs of individual neurons normalized activity to the increasing force (100% for the maximum force applied before withdrawal) and mean (bold line), for the linearly Activated (left) and the other sensory non linearly responding neurons to the force applied (right). b. average normalized activity ± SEM response to a mechanical paw withdrawal (-2.5 to 2.5s) for the the linear (n=37) (top) and non linear (n=15) cell populations responding to the force applied (bottom).

**Figure S5 : Spinal Encoding of contralateral sensory stimuli.** a. Sankey plot illustrating the neuronal dynamics across both dorsal horns (n = 154) in response to an ipsilateral and contralateral paw stimulation (mechanical or thermal) for all categories (Chi²: df= 19.05, 6, p<0.0041). b. top: heatmap of a pinch zscore responses of the dorsal horn units (3 mice, n=73) in anesthetized condition, from ipsi (left) and contralateral (right) stimulations. all units are aligned to the ipsilateral pinch. bottom : mean of activated units to ispilateral pinches (left) , and their response to a contralateral stilmulation (right) (n = 9, z-score ± SEM, bin = 0.05 s, -2.5 to 2.5 s).
