## Supplementary figures and images for "Spinotrode: long-term intraspinal electrophysiological recordings to unravel dorsal horn neuron dynamics in behaving mice"

### Fig S1

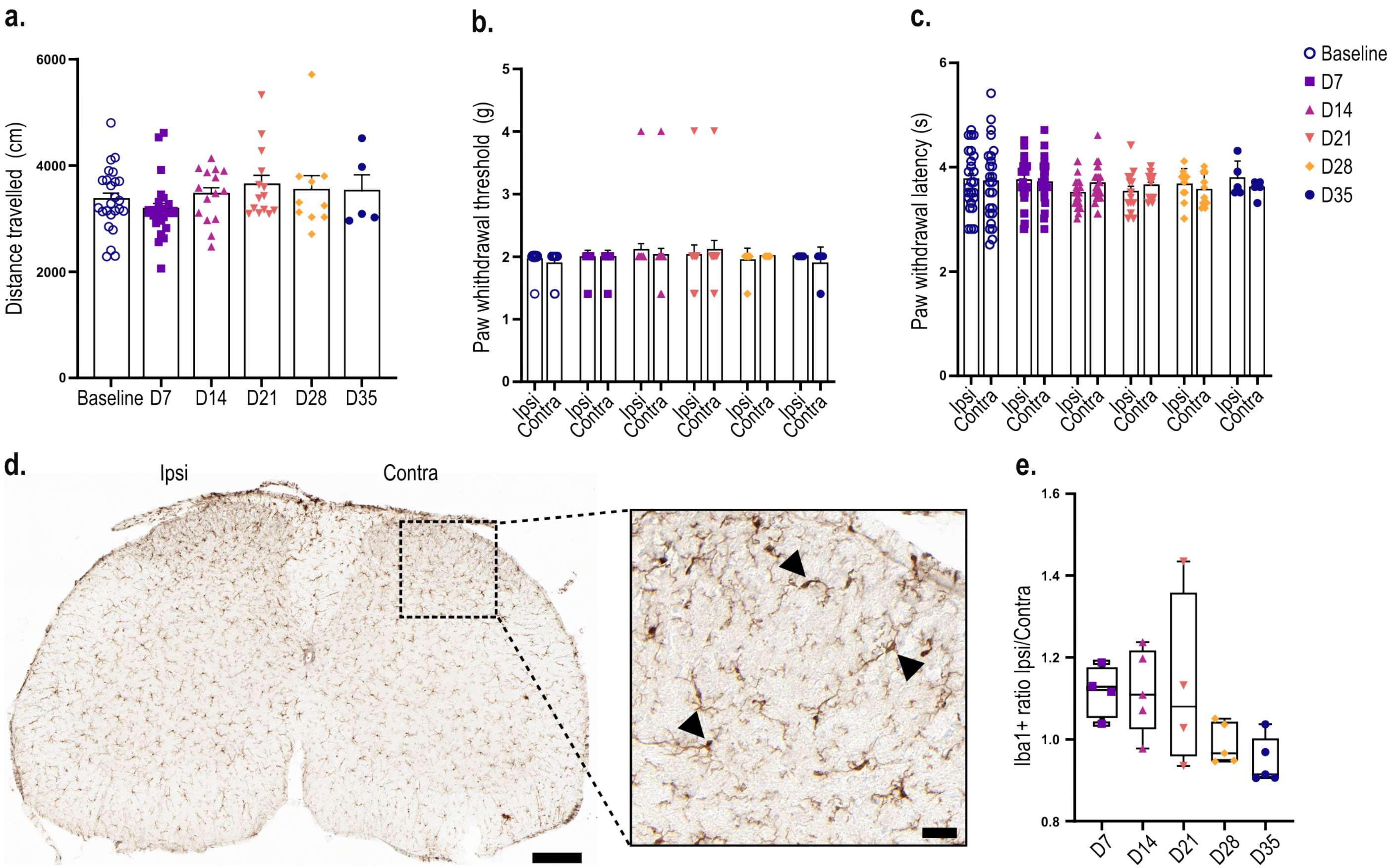

### Fig S2

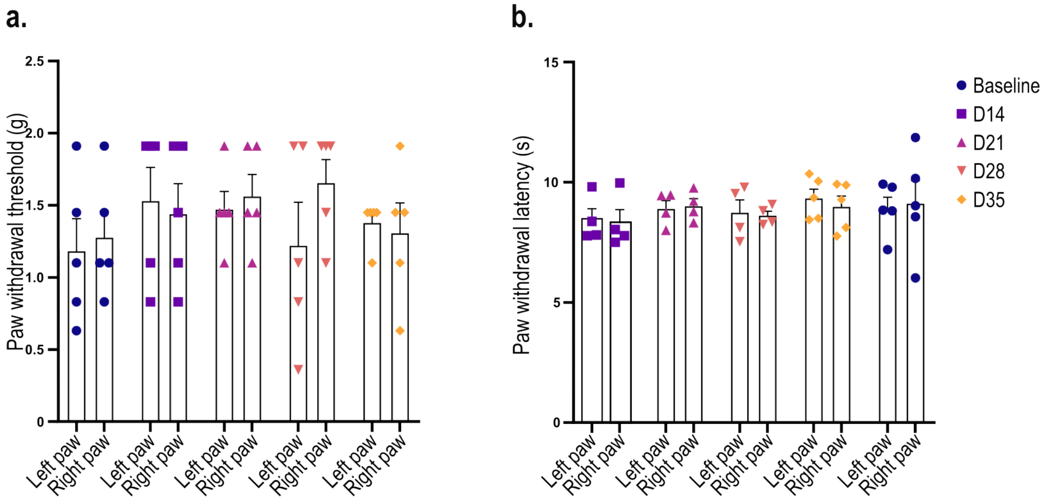

### Fig S3

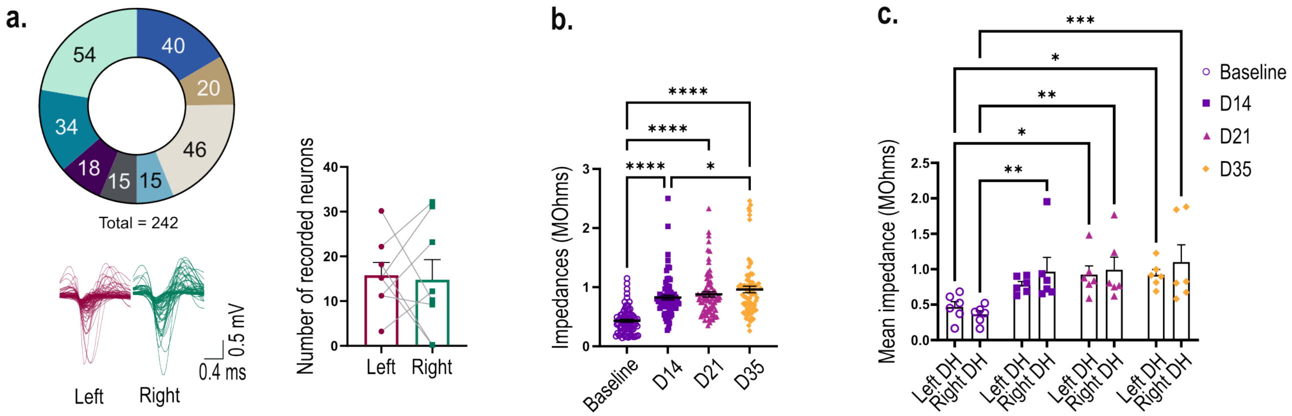

### Fig S4

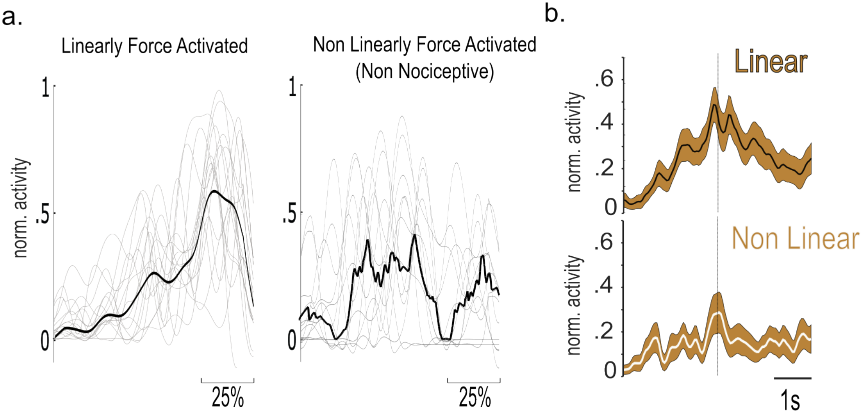

### Fig S5

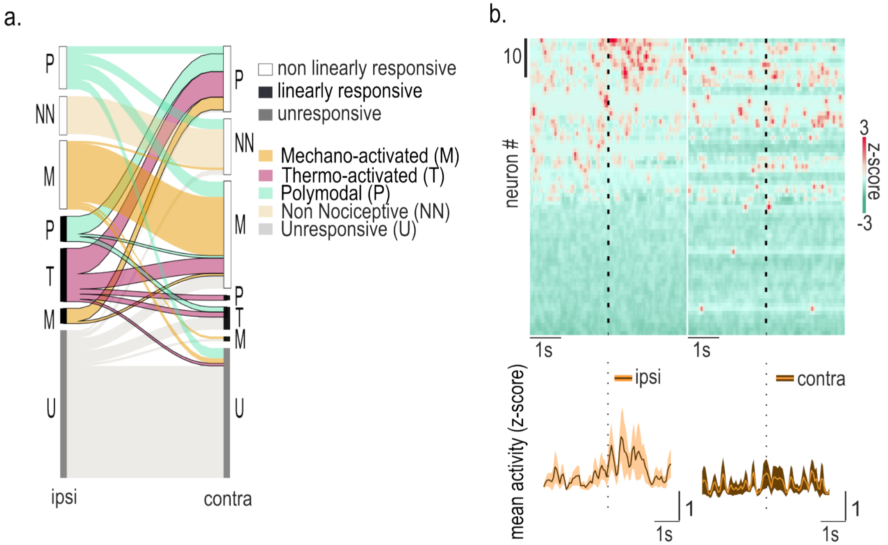
